# Ecological connectivity modelling for memory-constrained compute environments

**DOI:** 10.64898/2026.07.08.737333

**Authors:** Alex J. Southgate, Joanna Redihough, Thomas E. Woolley

## Abstract

Circuit theory has been successfully applied to ecological connectivity modelling, notably via the Circuitscape [1] software, which is typically run locally on a laptop, or on a dedicated server. However, for some applications, such web apps, automated pipelines, or high-throughput parallel analyses, backend infrastructure is often required. These systems can be costly, complex to operate, and introduce privacy and data governance concerns. Recent developments in WebAssembly (Wasm) now also allow fast C^**++**^ or Rust code to be run directly in a sandboxed browser environment. However, these are also contexts in which memory is limited. In order to make connectivity analysis feasible under memory constraints, we developed a Rust/WebAssembly toolset with a geospatial data pipeline and efficient, memory-optimised implementation of connectivity analysis. This approach may be useful for geospatial modelling software where raster sizes and memory are constrained, such as in the browser context. We also demonstrate that lower memory presents opportunities for parallelism, such as in the context of algorithms that require sampling, enabling previously constrained analyses. Our approach has the potential to reduce backend operational complexity, resource requirements, and overall cost.

## 1 Introduction

Connectivity analysis has found useful application in ecological modelling, population genetics, and conservation [1, 2, 3], including software such as Circuitscape [4], Omniscape [5] and R packages such as `gdistance` [6]. Tools employing circuit theory, such as Circuitscape, offer advantages beyond least-cost methods [1], which is available in a range of software and packages across different languages (e.g. Linkage Mapper [7], `pycda` [8], Ar-cGIS [9]). At the same time, these modelling approaches can require large compute resources, such as cloud instances or servers, which in turn require technical expertise to operate and maintain. As an example of scale, the Julia implementation of Circuitscape was previously benchmarked using 384 GB of RAM with a 437 million pixel input [10]. In the context of automated analyses, such as in bioinformatics workflows or web applications, batch cloud executors are often used (such as Google Cloud Batch) [11], and cost can scale with resource requirements.

In modern web applications, a front-end browser application is often paired with a backend server which performs more computationally expensive jobs. When scientific code is deployed into production web applications for practical use by customers (e.g. analysts), a backend is required. However, for many applications, maintaining this infrastructure is costly. One alternative is to offload computations browser-side. This can also be advantageous for data governance, as storage of personal data can be avoided if computation occurs directly in the user’s browser.

The problem with performing complex calculations in the browser is that browser code (i.e. JavaScript) execution is strictly controlled for security purposes, which has poor performance and filesystem interaction. To resolve this problem, WebAssembly [12] was introduced in 2017, allowing fast C^**++**^ or Rust code to be run in a sandboxed browser environment. It has previously been argued that WebAssembly is poised to transform scientific computing [13]. In addition, web applications may integrate complex user controls (e.g. modelling controls directly on a map) that are time consuming to complete using scientific scripts, requiring expertise not always available to users outside of academia.

WebAssembly has found use in several scientific projects, including: Pyodide, a CPython port, supporting installs of NumPy and SciPy, amongst other packages [14]; WebR [15], a WebAssembly compilation of the R language; ITK-Wasm, for spatial/image analysis [16]. Application domains include software for mapping pipeline corrosion [17], viral genomics [18], or cancer genome visualisation [19].

This paper introduces a Rust library and WebAssembly bindings to allow connectivity analysis in contexts where memory is constrained or costly, such as geospatial web applications. We demonstrate reasonable performance provided input images fit within browser sandbox memory limits, which has potential use for deployed scientific applications.

## 2 Methodology

### 2.1 Connectivity model

We use the methodology introduced by [3] to describe connectivity with circuit theory. First, Ohm’s law [20, Ch. 2] (1) specifies the relationship between voltage *V* , current *I*, and resistance *R* as *V* = *IR*. In the case of ecological connectivity, *I* is the movement of the migrating population instead of electrical current, resistance describes how hard it is for the population to move (consider movement over a rocky beach versus flat grassland), and voltage comparable to the pressure organisms experience to leave or enter a region (behaviorally or otherwise). Or, using conductance *C* = 1*/R*,

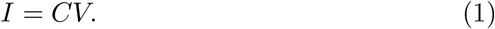

Secondly, for a given node (pixel), the conservation equation (Kirchhoff’s equation, see [20, Ch. 2]) means that movement in to a pixel is equal to movement out of the pixel. This does treat the modelled species as a conserved quantity and the simplest approach ignores factors such as death. Taking *S*_*i*_ as a source at node *i* (a point at which the organism emerges,such as a roost), with neighboring pixels *j* ∈ *N* (*i*):

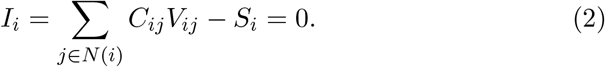

Given (1) and (2), only voltage differences, *V*_*ij*_, are unknown. Voltage, as a potential, can be assumed to be a scalar quantity for a single node up to some constant, which leads to a system of equations:

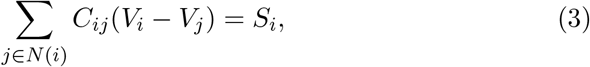

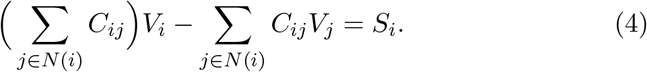

Equations (3) and (4) can be written in the form

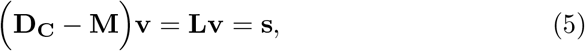

where the matrix **L** is the graph Laplacian on *N*. Equation (5) is an example of a discrete Poisson problem [21, Ch. 4].

For a ground (sink) node *j*, zero voltage Dirichlet boundary conditions can be enforced during solving by setting the corresponding voltage **v**_*j*_ to zero and skipping the corresponding rows in the calculation. For Robin-type conditions for ground, instead of a subset of pixels representing ground itself, the specified pixels are connected to an external ground (see the similar approach used in [22]). In this system, we use the modified Laplacian **L**_*shunt*_. On the diagonal, for a given pixel at index *i*, we add the shunt conductance *a*_*i*_, representing conductance from that pixel to the external ground. In this system, the voltage is not zero as in the Dirichlet case, and the external ground voltage is *g*_*i*_. The modified Laplacian has diagonals

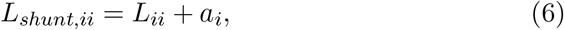

and the modified vector **b**_*shunt*_ has entries

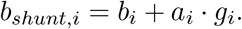

Notice from the units of each side that *a*_*i*_ is a conductance, *a*_*i*_ · *g*_*i*_ a current, *g*_*i*_ a voltage, and *a*_*i*_(*g*_*i*_ − *v*_*i*_) the current through the shunt.

### 2.2 Modes

The simplest run mode is for a source and sink pair, equivalent to a conditional random walk (paths of those that travel from source to sink). Circuitscape offers an ‘advanced’ mode, where a raster specifying sources and grounds can be provided, and a ‘pairwise’ mode [10], for which users can specify 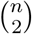 pairwise solves from pairs of *n* nodes, or *n* × *m* simulations between *n* sources and *m* sinks. In Omniscape, a circular moving window is used with a central ground, computed across the entire landscape, and combined [5].

Each approach has different assumptions and interpretation. We implement an equivalent to the ‘advanced’ mode, with 0*V* ground pixels or pixels connected to ground via shunt, with fixed source injection.

### 2.3 Solver

In Circuitscape, conjugate gradient with algebraic multigrid or a Cholesky solver is used via optimised libraries that are not designed for the Wasm context. In order to keep the Wasm binary small, we implement a conjugate gradient (CG) solver and compare several preconditioners (see below).

A good description of the CG method can be found in [23] or [21, Ch. 4]. Since we implement it manually for Wasm, a brief explanation follows. We solve the system

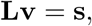

which is equivalent to minimising the energy function

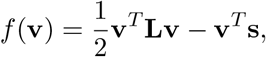

since, when taking the negative gradient, we get the residual error for our system,

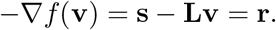

For an iterative solver, at iteration *k*, the residual **r**_*k*_ is computed. Stepping a distance *α*_*k*_ in the direction **r**_*k*_ at step *k* gives us the new guess

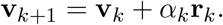

Since *f* is a quadratic function, it is convex given a symmetric **L** [24, Ch. 3.1.4]. Taking the gradient and choosing an optimal step size *α*_*k*_ achieves the global minimum for a slice of *f* (**v**), in the direction **r**_*k*_. Using this approach, the next step will be orthogonal to the previous one, since if it were not, the previous one would not have been the furthest (best) possible step. In steepest descent, to get the best step, applying calculus to the energy function it can be shown (see [21, Ch. 4]) that

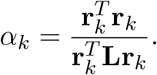

However, stepping maximally in a sequence of steps can be inefficient and lead to zig-zag steps, because each step is only orthogonal to the last step, not necessarily to all previous steps. Instead, the CG method guarantees orthogonality of residuals for all previous steps by enforcing conjugacy of step directions. Contrary to steepest descent, each direction requires information from the previous direction.

For CG, the conjugate direction **p**_*k*+1_ is taken to be

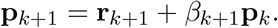

This is similar to steepest descent but has an additional term depending on the previous direction. **p**_0_ is initialized to **r**_0_. As before, once the step direction is taken, the position is updated as

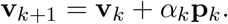

In the case of CG, *α* and *β* are [21, Ch. 4.3]:

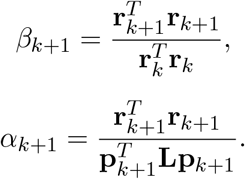

#### 2.3.1 Preconditioning

As described in [21, Ch. 4.3.4], convergence rate depends on condition number. For nonsingular symmetric matrices, the *κ*_2_ condition number is *κ*_2_(**L**) = ||**L**||_2_||**L**^−1^||_2_, where ||**L**||_2_ is the 2-norm (see [21, Ch A.3.2]). For symmetric positive-definite matrices with eigenvalues *λ*_*j*_, *κ*_2_(**L**) = max_*j*_ |*λ*_*j*_|*/* min_*j*_ |*λ*_*j*_|. For ill-conditioned matrices, where *κ*_2_ ≫ 1, steepest descent is slow. To reduce the condition number and speed up convergence, we solve a transformed system

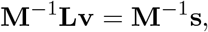

where **M**^−1^ approximates **L**^−1^. When **M**^−1^ = **L**^−1^, the result is the identity matrix, with optimal condition number of 1. We cannot compute **M**^−1^ directly, since this is computationally infeasible for large matrices, and if we could, we could solve our original system directly. Instead, we choose an easily invertible **M** that approximates **L**.

We compare two preconditioners: a simple diagonal preconditioner with **M** = diag **L**, also known as a Jacobi preconditioner, which can be useful for some data [25], smoothing high-frequency error components quickly, but low-frequency components slowly [21, Ch. 4.6]; a geometric multigrid (GMG) preconditioner, with bilinear interpolation, which is more complex but converges faster [26]. We implement these methods directly to reduce Wasm dependencies and binary footprint.

Multigrid preconditioners are more complex than the Jacobi preconditioner. After smoothing high-frequency error features using a few Jacobi iterations, the remaining low-frequency components are then smoothed using a coarse grid. A system **L**_*l*_**e**_*l*_ = −**r**_*l*_ is setup, with **L**_*l*_ and **r**_*l*_ coarse approximations of **L** and **r**, and **e**_*l*_ an approximation for the error **e** = **v** − **L**^−**1**^**s** in **v** [21, Ch. 4.6]. The subscript *l* refers to the level, or the number of times this process is repeated recursively. As the final output, the **e**_0_ is often referred to as the preconditioned residual.

Construction of coarser systems is recursive, so that the error in the second level refers to the error in the error from the first level. As such, at each level within the V-cycle, the residuals of the system are not the same as the outer-layer CG residuals, but instead are residuals of the *l*th system. Once this system is solved, interpolation is used to return to the fine grid, which works for low-frequency features with high-frequency features already having been smoothed out. Finally, smoothing is performed again to eliminate interpolation artifacts. If this process is applied recursively, one outer loop is known as a ‘V-cycle’ [21, 27, Ch. 4.6]; at the lowest resolution, the system is solved directly.

Overall, one cycle performs the application of matrix **M**^−1^. For the CG update, the preconditioned residual **e**_**fine**_ is used to update **p** as [28, Ch. 9.2]

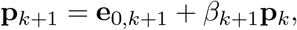

with

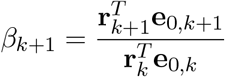

#### 2.3.2 Memory optimisation

We implemented a memory-optimised mode for the solver in order to trade time for memory efficiency in the browser context where memory constraints can be more important. The first optimisation we make is to take a hybrid approach to matrix storage. The largest data structure during run-time is the fine Laplacian, with off diagonal entries

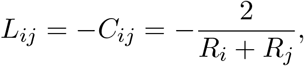

and diagonal entries

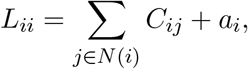

where *a*_*i*_ is the shunt, as in 6.

In the explicit representation, this Laplacian can be pre-computed before the CG algorithm. This is straightforward to replace with an implicit representation that computes entries lazily from the input conductance values with an additional averaging operation per access.

The level-1 Laplacian can also be made implicit to avoid storage. As described, the level-1 Laplacian is:

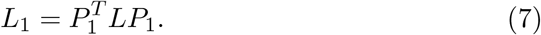

The number of non-zero terms in computing each element of *L*_1_ depends on the operator used. For a 2D raster grid, each row and column of *L* has at most 5 non-zero entries. For a 9-point bilinear prolongation, each row of 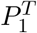may have 9 non-zero entries. Unlike with the fine matrix, computing each entry of *L*_1_ can be costly, resulting in a tradeoff between runtime and memory.

For pre- and post-weighted Jacobi smoothing, we use separate parameters for the number of smoothing iterations *η* and weighting parameter *ω*, since smoothing the level-0 and level-1 implicit matrices is more costly than smoothing the explicit coarse laplacians.

For a number of our tested scenarios, 32-bit mode for Circuitscape and our own 32-bit implementation failed to converge. As such, we additionally implemented a mixed-precision implementation using 64-bit intermediates and a 64-bit voltage vector to improve convergence while minimizing memory usage. We assessed this mixed-precision mode with our full 64-bit implementation.

### 2.4 Test datasets and benchmarking

Two datasets were used for assessment, covering two regions. Firstly, a test region corresponding to a 1 km^2^ region centered at 286600E, 79050N (EPSG:27700 easting & northing coordinate system) was used for time and memory benchmarks, referred to here as the Chudleigh dataset. For the three-scenario comparison, a region centered at 290959E, 55599N was used, referred to as the Brixham dataset. These regions include building, road, and river features. Vector data was obtained from the UK Ordnance Survey open data [29]. Digital surface model (DSM) and digital terrain model (DTM) LIDAR data was obtained from the UK Environment Agency [30]. We used an additive model for landscape resistance as described in [31].

For the Chudleigh resistance map, we use constant resistance parameters for rasterised vector features: 5 for roads (3 meter buffer), 500 for buildings, 0.5 for rivers (4 meter buffer). For raster landscape feature resistance, we took the feature height as resistance and multiplied by 100. These parameters were arbitrary and chosen to introduce variation into the test resistance map inputted into current calculation. In practice, these parameters can be set freely.

For comparison of our low-memory approach with Circuitscape v6.1.0, we used three scenarios from three realistic resistance maps, spanning maps rich in both low- and high-frequency variation. We used a final residual tolerance of 1 × 10^−4^ for both 64-bit implementations and 1 × 10^−3^ for our mixed-precision implementation. For the three scenarios, parameters for features such as roads and point lights were taken from [31]. These scenarios are shown in 4.

Reduced memory consumption raises the potential for greater parallelism with fewer compute resources. To demonstrate an application of this, we applied our method to assessment of sensitivity of current maps to changes in underlying resistance map parameterisation. The third scenario of the Brixham dataset was used. In this analysis, 100 random current maps were solved. Each current map used the same resistance parameters as previously,except for the light maximum resistance value, which was randomly sampled as 10^*U*^ with uniformly sampled *U* ∈ [3, 8], in order to sample across orders of magnitude.

Benchmarking was undertaken on a laptop with an AMD Ryzen 7 5700U, 16G of RAM, Linux 6.8.0-51-generic, and using Google Chrome 150.0.7871.46. Benchmarking code was wrapped in a ReactJS [32] applet which can be run locally and accessed via the code repository https://github.com/asouthgate/wasm-connectivity.

### 2.5 Code

Code can be obtained at https://github.com/asouthgate/wasm-connectivity.Optimisation codes were written in Rust and compiled to Wasm using `wasm-pac`k [33] and `wasm-bindgen` [34]. This code contains several simple examples. All results can be easily reproduced by running the ReactJS [32] development server, clicking to the benchmark panel, running the benchmarks, and downloading the results.

## 3 Results

Figure 1 shows time and memory benchmarks as a function of raster size for the explicit MGCG, memory-optimised MGCG, and low-memory mixed-precision MGCG modes, tested on the Chudleigh example dataset. For the low-memory mode, run-time increased modestly for a large reduction in memory. For the 2000 × 2000 rasters, the low-memory and low-memory mixed-precision implementation used under 300 MB, compared to 800 MB used in the explicit mode. This trade-off was reflected in runtime, where solving the largest rasters took approximately 3× longer. For the browser context, this trade-off can enable workloads not practically possible otherwise, such as for high-resolution LIDAR data at 1 metre per pixel, covering several kilometres. Figure 2 shows a side-by-side comparison of a rescaled Circuitscape current map produced from the same resistance map input with a zero-volt ground, showing equivalent results (max absolute difference of*<* 10^−5^).

**Figure 1:**
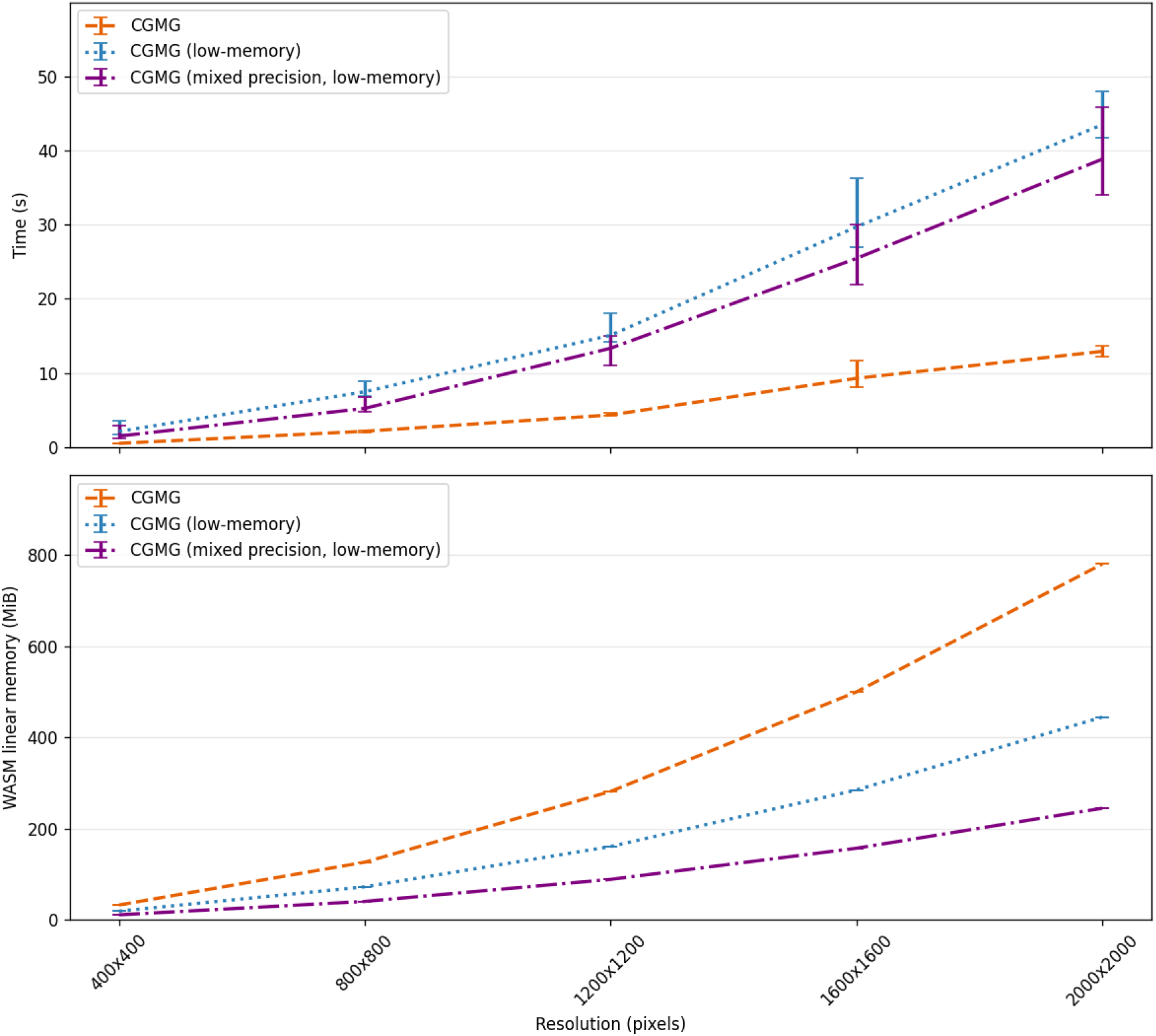
Time and memory comparison across resolutions for each mode. Bars represent sample range, with the mean as lines. The number of iterations at each resolution were 12, 12, 11, and 13 for explicit and low-mem versions, and 12, 12, 10 and 9 for low-mem mixed-precision.

**Figure 2:**
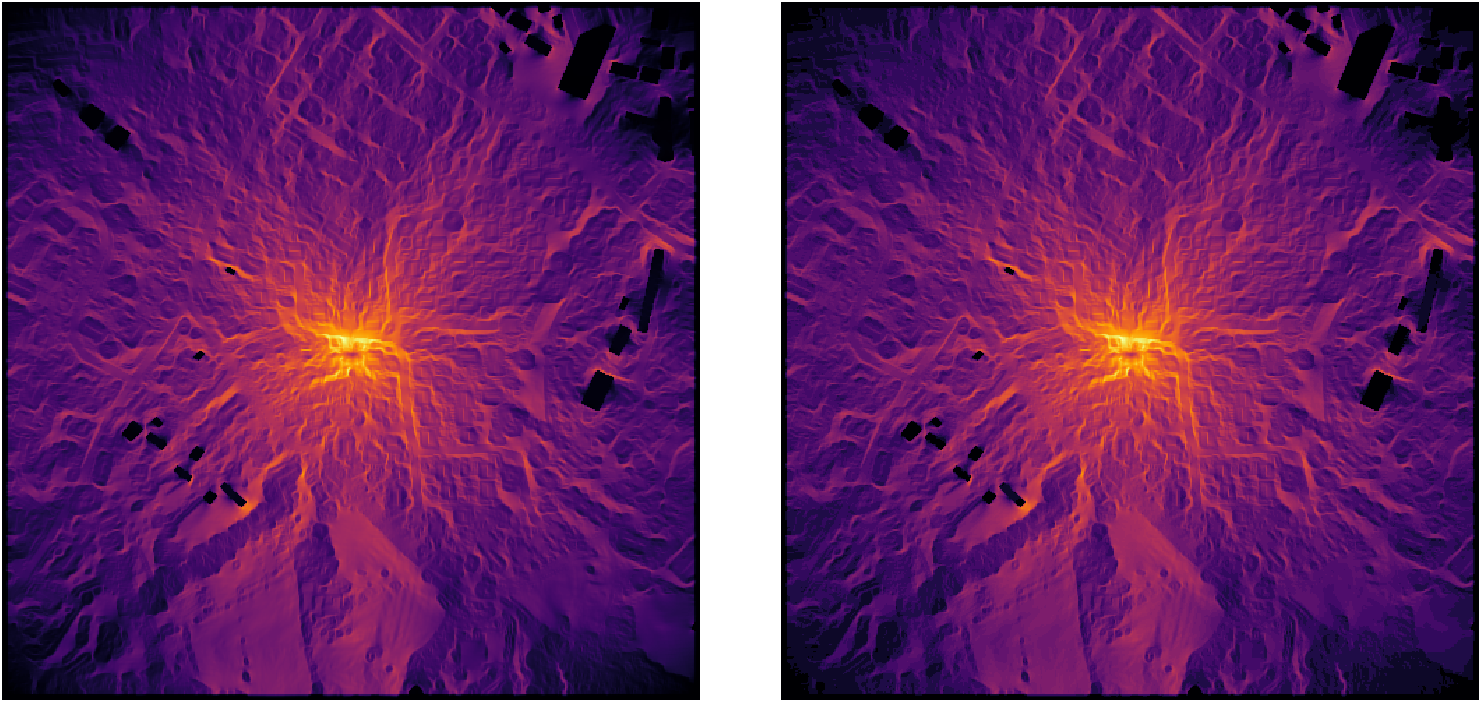
Side-by-side comparison showing log rescaled current maps produced by Circuitscape (left) and our low-memory Rust solver (right), to show consistency. Current maps were rescaled to the range [1, 100]. Max absolute difference *<* 10^−5^ rescaled units.

Since performance can vary depending on input raster, we additionally assessed performance across a range of scenarios using the Brixham dataset. As demonstrated in Figure 3, our approach in 64-bit mode results in an approximately 7-fold reduction in memory when compared with Circuitscape,from 3.5 GB to 500 MB for 2000×2000 raster images, with comparable runtime. Our mixed-precision mode used under 300 MB, while 32-bit modes for both Circuitscape and out implementation failed to converge. For each of the scenarios, our 64-bit low-memory implementation required 12, 9, and 27 iterations respectively. Figure 4 shows these scenarios, each of which is non-trivial. Each scenario is derived from DSM, DTM, and open vector data covering a test region. The third scenario (bottom row) features several orders of magnitude change in resistance values and discontinuous features corresponding to roads and point lights. Despite these features, mean run-time was under 70 seconds. Although the number of iterations may increase for other regions and resistance parameterisations, we expect similar run-times across geographic regions.

**Figure 3:**
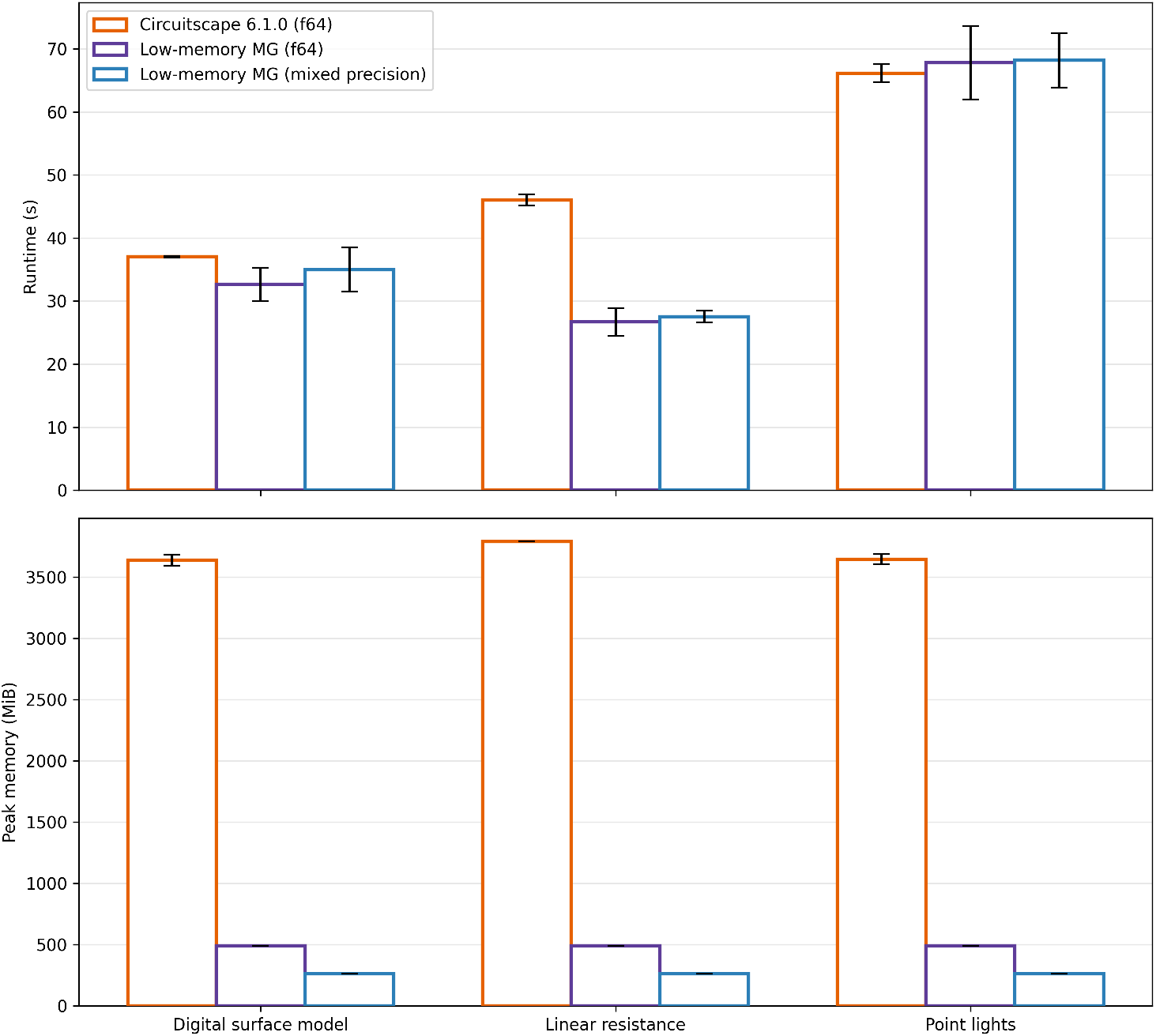
Runtime and memory comparison between Circuitscape v6.1.0 and our low-memory implementations. For all three scenarios, there was an approximately 7-fold reduction in memory, from 3.5 GB to 500 MB, with comparable runtimes. For mixed-precision, memory was 10× lower, although this would be expected to less compared to 32-bit Circuitscape, which did not converge for the given input scenarios.

**Figure 4:**
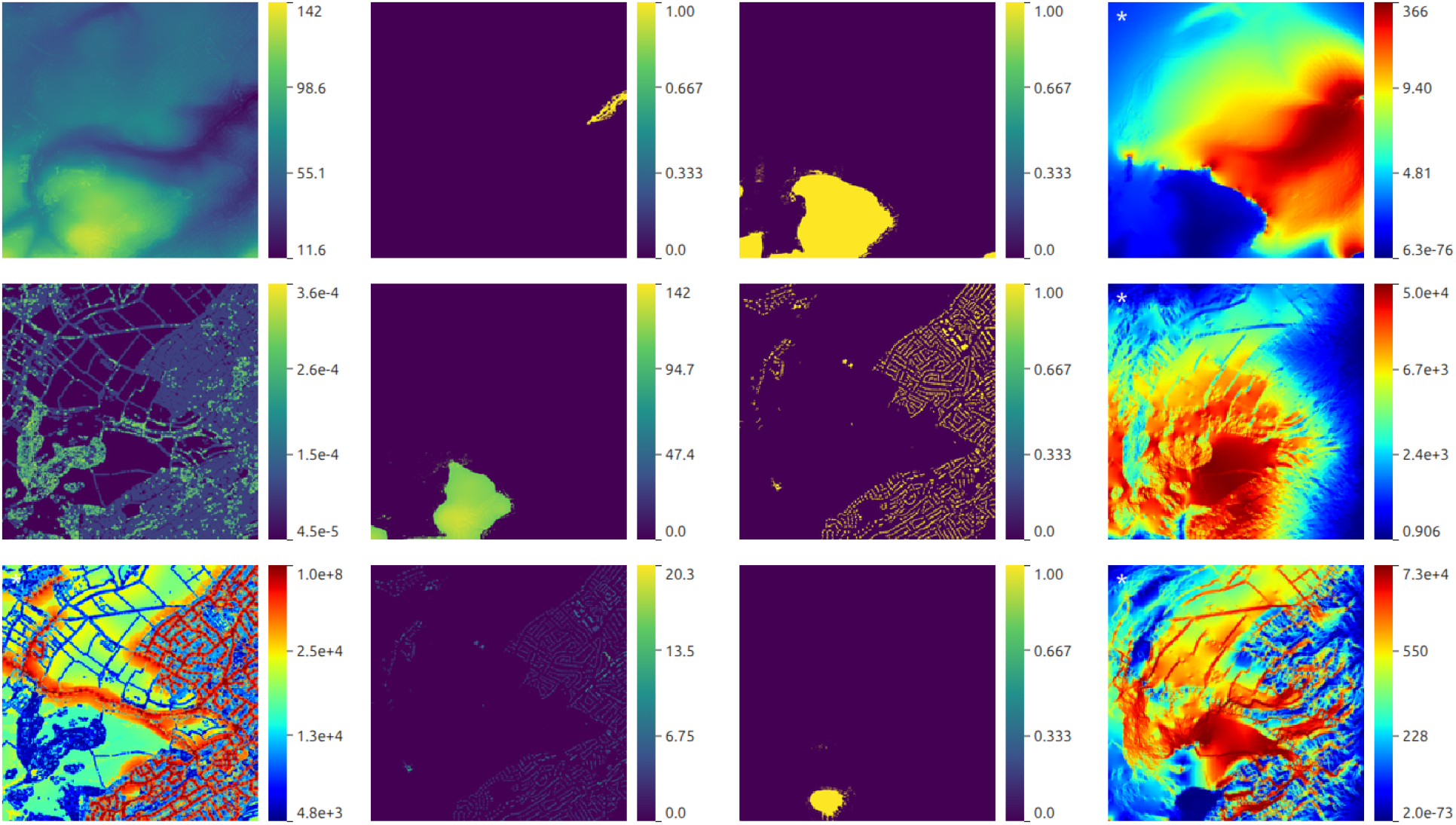
Three scenarios used for 2000×2000 rasters tests. Each column represents resistance, source, ground, and computed current maps. The point-light resistance (bottom left) and current maps (right column) are quantile colormapped (white asterisk) for improved visualisation. The scenarios include varying features. The third contains resistance values spanning several orders of magnitude, with sharp discontinuities across the landscape.

Reduced memory opens up the possibility for greater parallelism and further use cases. We demonstrated this by undertaking an assessment of current sensitivity to resistance map parameterisation. The Brixham point-light scenario was used again for this analysis. Although optimal parameters for the resistance maps used for this were derived from [31], we wanted to additionally assess sensitivity to parameter values. In this parameterisation, maximum light resistance exceeds all other resistances sources by several orders of magnitude, at 10^8^. We ran 100 concurrent analyses with randomly sampled maximum light resistance between 10^3^ and 10^8^. Figure 5 shows the minimum (1 × 10^3^), median (9.3 × 10^5^), and maximum (9.6 × 10^7^) values sampled. Across these resistance values, several key boundaries only become visible at the higher values (c). In (d), these boundaries also correspond to regions which high standard deviation across the samples. Some qualitative features are relatively stable across resistance values. For example, lines corresponding to hedgerows appear in all three current maps with varying intensity. However, in (d), features in red are not consistent across the maps, which largely correspond to regions of very low current around lamp features. This analysis demonstrates that it is not simply sufficient to have high resistance values, and that, for example, 10^8^ produces a qualitatively different map from 10^6^. This analysis completed in 17 minutes on a laptop, which would not be feasible if each analysis took several gigabytes of memory.

**Figure 5:**
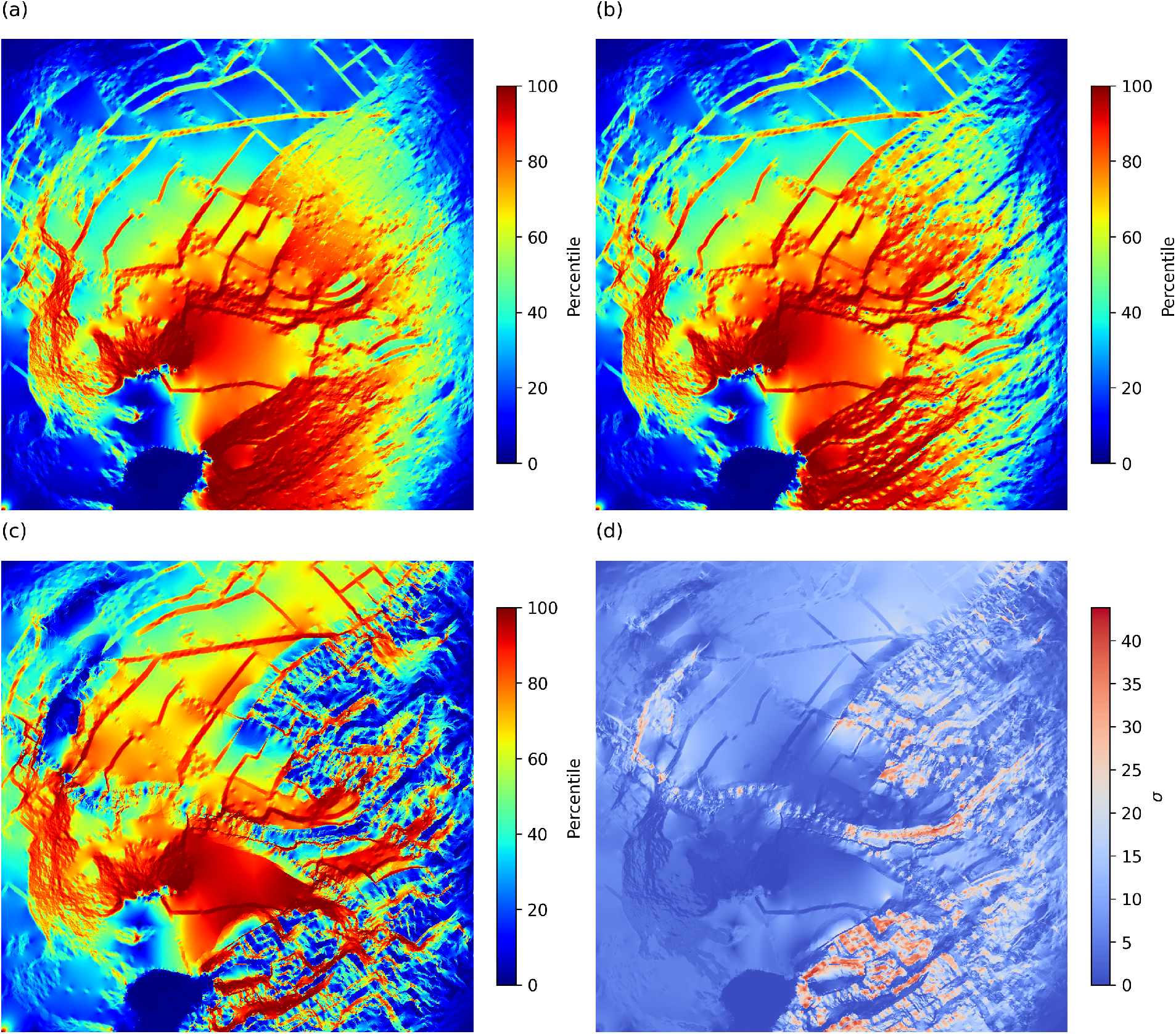
Point-light scenario with sampled point-light maximum resistance values between 10^3^ and 10^8^. Panels (a), (b), and (c) show single-sample current maps with sampled minimum (1.0 × 10^3^), median (9.3 × 10^5^), and maximum (9.6×10^7^). As can be seen in these panels, a qualitative difference is observed between the median and maximum, implying that a resistance of 9.5 × 10^5^ is insufficient to capture the barriers that form with a resistance of 9.6 × 10^7^. (d) shows standard deviation of quantile values across samples.

## 4 Discussion

We present a Wasm/Rust connectivity library for interactive geospatial edge-computing in memory-constrained environments. Our memory-efficient MGCG solver demonstrates comparable runtimes to Circuitscape with lower memory. This method can be integrated into geospatial web applications, allowing connectivity modelling in the resource-limited browser Wasm context. Our approach can also reduce the required backend server resources, as well as downstream concerns such as data governance complexity.

Another promising opportunity of our approach is that of increase parallelism. Algorithms that require repeatedly computing current maps from sampled resistance maps, such as for characterising uncertainty, may be more feasible. We used 16 threads to compute 100 current maps in 17 minutes on a laptop. This could widen the connectivity analyses possible on local resources, and as shown in our results, previous methods requiring multiple gigabytes per analysis would likely have taken several times longer given constrained memory. In principle, using 16 threads, thousands of samples could be computed on a laptop over the course of hours.

Different solvers have different performance on different input matrices. While this implementation gives reasonable performance for the tested data across several realistic scenarios, it is likely to vary with different rasters depending on the type of features they contain. For certain resistance maps, operator-dependent restriction, such as Alcouffe’s method [27] could be used. Limitations of the Wasm environment are primarily constraints in memory, CPU, and binary size. Traditionally, scientific software relies on highly optimised libraries such as BLAS [35]. In principle, this is possible for Wasm, but it is not straightforward. Projects like Pyodide [14] demonstrate the work required for compiling existing packages into Wasm executables. In addition, for front-end web applications, it is often important that data transfer is minimised to enable responsive pages.

## 5 LLM disclaimer

This paper was written by hand, without LLM generation. Code and research was developed with LLM assistance.

## 6 Contributions

Alex Southgate: conceptualization; problem formulation; solver and geospatial pipeline implementation; low-memory optimisations; paper writing; test-ing. Joanna Redihough: conceptualization, especially with respect to utility of frontend integration; front-end code; paper editing. Thomas E. Woolley: paper editing; conceptualization of improvements to benchmarking.

## Notes

### Competing Interest Statement

The authors have declared no competing interest.

### Summary of Updates

- Added another analysis demonstrating improved parallelism that can be used for evaluating sensitivity to resistance parameters - Added a new figure - Improved methodological descriptions - Edited text

